# ClusToRa: A niche-centric framework for identifying structural recruitment and infiltration in spatial omics

**DOI:** 10.64898/2026.05.23.727398

**Authors:** John Maringa Githaka, E. Paul Lerner

## Abstract

Spatial omics maps cellular landscapes, yet current tools might conflate stochastic proximity with organized niches. We present ClusToRa (Cluster-to-Randomization), a framework that identifies high-density cellular territories and quantifies cell-type recruitment using a fixed-position null model. Benchmarked against graph-based neighborhood-enrichment and point-pattern statistics, ClusToRa reduced false-positive enrichment in simulations and resolved core-vs-boundary interactions. Applied to cirrhotic MASH liver, ClusToRa identifies stellate-cell territories with immune/endothelial infiltration and stress-, Notch-, and PPARα-associated programs, providing a niche-centric framework for distinguishing structural cellular infiltration from boundary adjacency or density-driven colocalization.

## Main

Tissue function emerges from spatially organized cellular interactions. Graph-based neighborhood analyses capture local adjacency but are sensitive to global density variation and lack geometric resolution to distinguish incidental proximity from structured infiltration^1,2^. Label-randomization approaches ignore spatial niche boundaries, while classical point-pattern statistics such as Ripley’s functions global summaries but do not localize interaction domains and assume isotropy, limiting their applicability to irregular tissue geometries^3,4^.

We developed **ClusToRa** (Cluster-to-Randomization) to explicitly model spatial territories and quantify cell-type recruitment (Fig.1a). ClusToRa identifies cellular domains using DBSCAN with an automatically determined neighborhood radius (ε), followed by a fixed-position null model that preserves spatial coordinates while randomizing cell identities. This enables estimation of recruitment-specific Z-scores while reducing susceptibility to density-driven false-positives. To capture domain geometry, ClusToRa reconstructs cluster boundaries using α-shapes, enabling classification of interactions into **core infiltration** and **boundary envelopment**, and quantification via an infiltration–envelopment ratio. At the tissue scale, spatial niche organization is summarized using the Clark–Evans index (R), distinguishing aggregated, random, and dispersed distributions.

**Figure 1:**
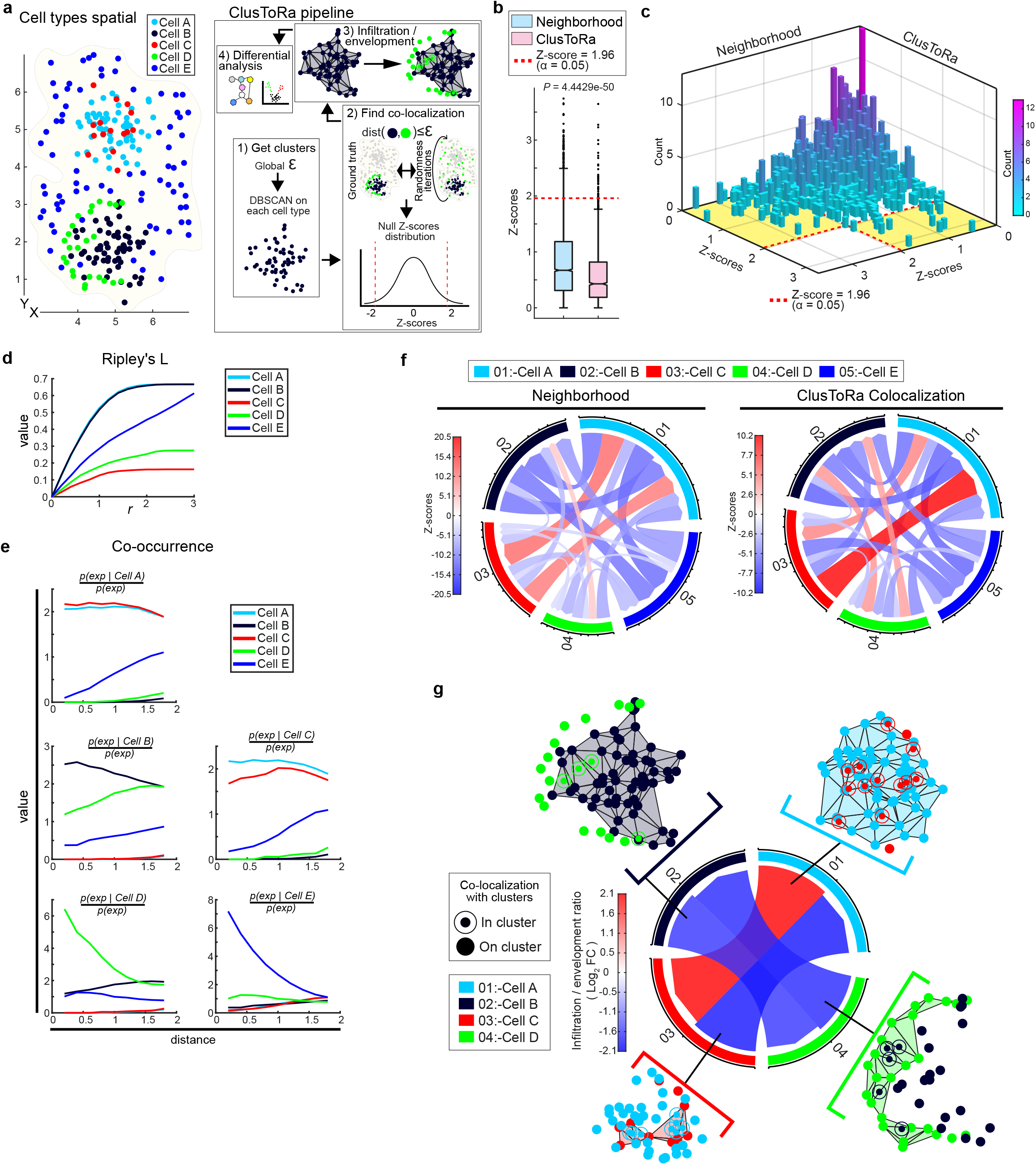
Development and benchmarking for ClusToRa. **a**, Simulated spatial distribution and ClusToRa workflow. Left: representative simulated cell types illustrating predefined infiltration and envelopment interaction patterns. Right: ClusToRa pipeline showing (1) density-based clustering of cellular territories using DBSCAN, (2) fixed-position null model for colocalization testing, (3) α-shape–based reconstruction of non-convex cluster boundaries with classification of interactions into core infiltration and boundary envelopment, and (4) differential analysis of those niches when omics data is available. **b**,**c**, Distribution of Z-scores under null simulations. Box plots (b) and histograms (c) show Z-score distributions from 100 null simulations (∼6,400 cells per simulation, 10 cell types in each). Neighborhood enrichment implemented in Squidpy produces a substantial fraction of false-positive interactions exceeding the nominal threshold (Z ≥ 1.96), whereas ClusToRa suppresses density-driven artifacts, with the majority of scores remaining below this threshold. **d**,**e**, Comparison with point-pattern statistics. Ripley’s L-function (d) and co-occurrence analysis (e) computed using Squidpy applied to simulated data in (a). These approaches capture global or distance-dependent spatial trends but fail to resolve discrete, non-convex interaction domains or distinguish infiltration from adjacency. **f**, Neighborhood enrichment and ClusToRa colocalization Z-scores chord diagrams. For A → B, B is evaluated relative to A-defined clusters . **g**, Chord diagram showing *log*_2_ infiltration–envelopment ratios: For A → B, red indicates B embedded within A-defined clusters; blue indicates B positioned alongside or outside A-defined boundaries.

Across simulated datasets with known ground truth, ClusToRa colocalization demonstrated improved specificity relative to neighborhood enrichment in Squidpy^5^, which produced significantly more false-positive enrichment under null conditions (Fig.1b,c). In contrast, ClusToRa Z-scores remained below 95% CI with minimal outliers, demonstrating improved specificity and robustness to density effects.

Compared to point-pattern statistics, Ripley’s L-function underestimated clustering in irregular domains, while co-occurrence analysis captured distance trends but failed to resolve discrete territories (Fig.1d,e). ClusToRa distinguished interaction patterns across complex geometries and identified relationships missed by graph-based approaches, with the infiltration–envelopment ratio uniquely resolving core versus boundary behaviors (Fig.1f,g).

To evaluate performance in a biological context, we applied ClusToRa to a spatial transcriptomics atlas of Metabolic dysfunction-associated steatohepatitis (MASH) cirrhotic liver tissue comprising control, high-BMI, and low-BMI cohorts (Fig.2a), derived from Tzouanas et al dataset^6^. MASH provides a stringent test case for ClusToRa because the fibrosis, inflammation, vascular remodelling, and hepatocellular stress creates dense, spatially heterogeneous tissue^7,8^. Prior single-cell and spatial transcriptomic studies identified stellate-macrophage colocalization and fibrosis-associated programs, and immune-mesenchymal signalling hubs^9–11^, but generally cannot resolve whether immune cells infiltrate stellate-defined territories or are simply restricted to their boundaries. ClusToRa addresses this by combining territory detection with fixed-position randomization and infiltration-versus-envelopment analysis.

**Figure 2:**
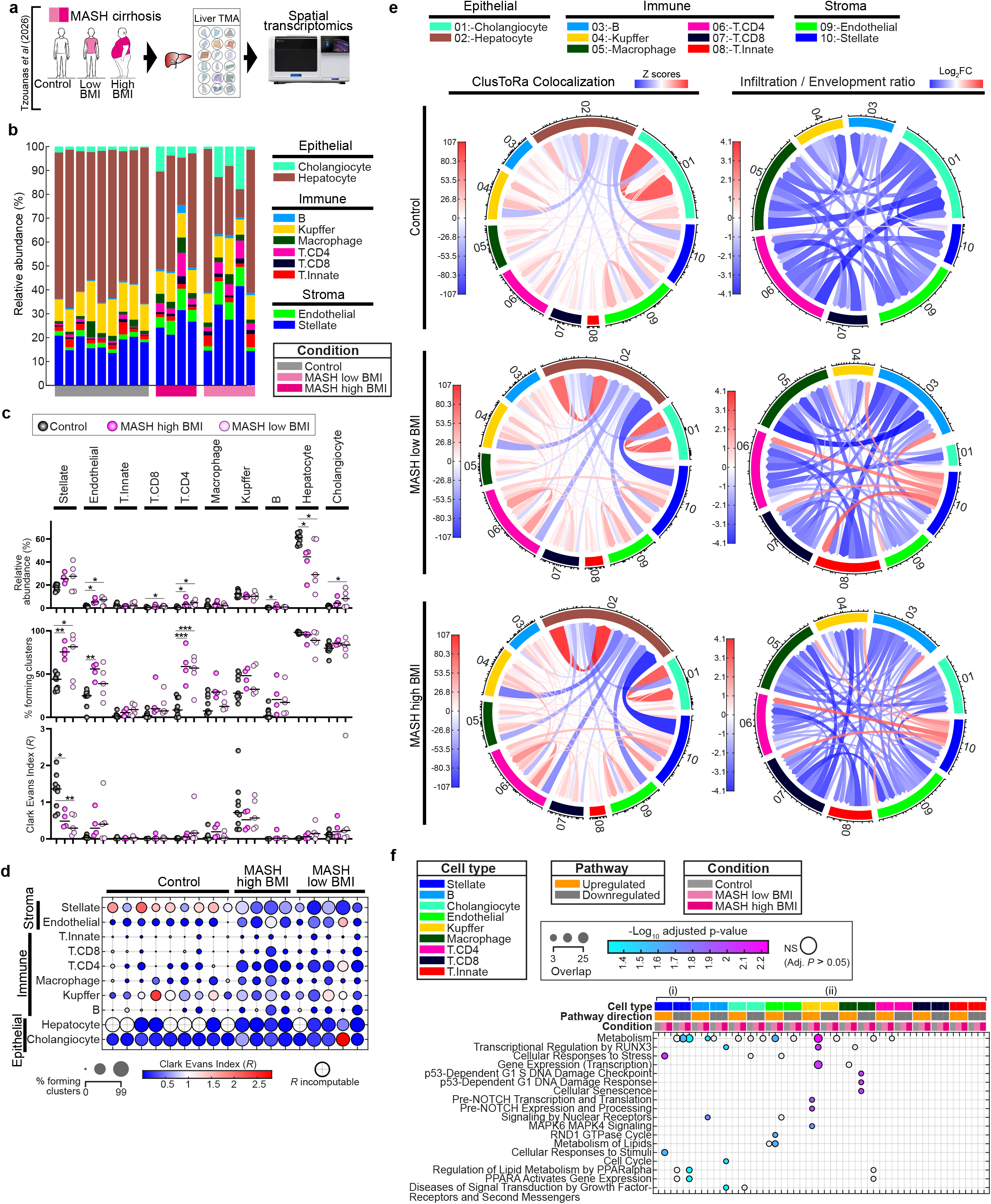
ClusToRa identifies stellate-centered niches as hubs of immune recruitment in MASH. **a**, Spatial transcriptomics dataset comprising control, high-BMI MASH, and low-BMI MASH cirrhotic liver tissue. **b**, Cell type relative abundance. **c**, Abundance and clustering across conditions. Significance assessed by one-way ANOVA followed by Tukey’s post hoc test (*P*<0.05, \**P*<0.01, \*\**P*<0.001). **d**, Cell type cluster distribution summary showing the proportion of cells forming spatial clusters (circle size) and their centroid Clark–Evans index (R; color scale). Increased clustering of endothelial, hepatic stellate, and CD4^+^ T-cells is observed in MASH relative to controls. Stellate territories exhibit aggregated spatial organization (R < 1) in MASH compared to dispersed patterns (R > 1) in controls. Cell types with fewer than three clusters were excluded from R calculation. **e**, ClusToRa colocalization and infiltration– envelopment networks identify stellate-centered immune/endothelial infiltration in MASH. **f**, Gene enrichment analysis of stellate-associated niches (Reactome pathways analysis via Enrichr^15^) for (i) clustered stellate cells versus non-clustered stellate cells and (ii) cells colocalizing with stellate territories versus non-colocalizing cells. Pathways related to metabolism and cellular stress responses are enriched, consistent with chronic metabolic adaptation. Colors indicate pathway directionality and sample group.

ClusToRa revealed disease-associated niche reorganization, including increased clustering of endothelial, hepatic stellate, and CD4+ T cells in MASH (Fig.2b,c). Stellate territories shifted from dispersed organization in controls (R>1) to aggregated distributions in MASH (R<1) (Fig.2d), demonstrating how territory-level analysis can resolve emergent multicellular architectural remodeling not captured by pairwise adjacency metrics alone. Notably, immune-stellate interactions were dominated by **core infiltration** rather than peripheral adjacency (Fig.2e). B cells, Kupffer cells, macrophages, CD4^**+**^ and CD8^**+**^ T-cells, and endothelial cells consistently localized within stellate-defined territories, whereas innate T-cell infiltration was selectively observed in low-BMI MASH. These findings are consistent with known stellate-immune crosstalk in fibrotic liver disease^12,13^ and suggest that stellate-rich fibrotic regions function as spatial organizational hubs for immune and vascular recruitment.

Gene enrichment analysis linked these stellate-associated niches to stress responses, PPARα, and Notch-related pathways, consistent with chronic metabolic adaptation and fibrotic remodeling(Fig.2f). The selective enrichment of innate T-cell infiltration in low-BMI MASH may reflect BMI-stratified differences in fibrotic niche organization^14^, although this observation remains hypothesis-generating given the modest cohort size and advanced cirrhotic stage of the source dataset.

Limitations include reliance on published cirrhotic MASH tissue, with uncertain generalizability to earlier disease stages, dependence on upstream segmentation and cell-type annotation. ClusToRa is therefore intended to complement existing spatial frameworks by providing territory-centric analysis of structured colocalization, including resolution of core infiltration and boundary organization beyond aggregate neighborhood enrichment approaches.

## Online methods

### ClusToRa framework

ClusToRa identifies spatially coherent cellular territories and quantifies cell-type recruitment relative to these domains. Spatial coordinates and cell-type annotations were used as input.

Cellular territories were defined using Density-Based Spatial Clustering of Applications with Noise (DBSCAN). The neighborhood radius (ε) was determined using a k-distance elbow approach. Briefly, the distance to the k-th nearest neighbor (k = MinPts, set to 4) was computed for each cell and sorted in ascending order to generate the k-distance curve. A line connecting the first and last points of this curve was constructed, and the perpendicular distance from each point to this line was calculated. The point with the maximum distance was identified as the elbow, and the corresponding distance value was selected as ε.

Clusters were defined as contiguous regions of points satisfying the density criterion of DBSCAN, enabling detection of arbitrarily shaped, non-convex spatial domains while excluding sparse background noise.

To quantify colocalization between cell types, ClusToRa employs a fixed-position null model. For a given query cell type containing *n* cells, *n* spatial coordinates are randomly sampled from all positions, effectively permuting cell-type labels while preserving the underlying tissue architecture. This procedure was repeated for 1,000 permutations, and colocalization counts were recomputed for each iteration.

A cell was considered colocalizing with a cluster if it fell within the ε-neighborhood of any point belonging to that cluster, ensuring consistency between clustering and interaction quantification.

Z-scores were calculated as:

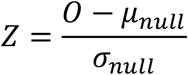

Where *O* is the observed proportion of colocalizing cells, and *μ*_*null*_ and *σ*_*null*_ are the mean and standard deviation of the null distribution, respectively. Statistical significance colocalization was defined using a two-sided threshold of *Z* ≥ 1.96 unless otherwise specified.

Two-sided empirical P-values were computed as:

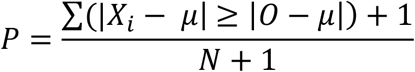

where X_*i*_ represents values from the null distribution, *μ* its mean, and *N* the number of permutations. The +1 correction was applied to avoid zero-valued P-estimates.

### Geometric reconstruction of territories

Cluster boundaries were reconstructed using α-shape complexes. The refinement parameter α was set proportional to the optimized ε, ensuring that the boundary topology remained consistent with the density-based clustering. This concave hull approach captures non-convex geometries and local voids that a standard convex hull would otherwise mask.

### Infiltration and envelopment classification

Colocalizing cells were categorized into distinct interaction modes based on their spatial relationship to the cluster’s geometric boundary. *Core Infiltration:* Cells residing within the interior of the α-shape volume, representing integration into the cluster core (Fig. 1g). *Peripheral Envelopment:* Cells situated within the local ε-neighborhood but remaining external to the α-hull, representing surface-level association (Fig. 1g). The infiltration–envelopment ratio was quantified as the *log*_2_ fold change between these two populations, where positive values indicate a preference for cluster infiltration.

### Tissue-scale spatial niche organization

Global spatial organization of cellular territories was quantified using the Clark–Evans index (R), computed from cluster centroids. Values of R < 1 indicate aggregation, R ≈ 1 randomness, and R > 1 dispersion. Only cell types with ≥3 clusters were included in Clark–Evans analysis.

### Simulation benchmarking

Simulated datasets were generated to model both structured and null spatial configurations. Null datasets were created by randomizing spatial relationships across 10 cell types (500–800 cells per type; mean total ∼6,400 cells) to establish a stochastic baseline. The structured coordinates incorporated predefined core infiltration and boundary interaction patterns, supplemented by random noise coordinates. ClusToRa’s performance was benchmarked against neighborhood enrichment and point-pattern statistics as implemented in Squidpy. To assess the false-positive rate, we evaluated the distribution of Z-scores under null conditions, where colocalization is purely stochastic and the spatial ground truth is known (Fig. 1b,c). We defined the Type I error rate as the proportion of tests yielding a *Z*-score > 1.96, corresponding to a 95% confidence level for significant colocalization.

### Spatial transcriptomics dataset

ClusToRa was applied to a published spatial transcriptomics atlas of MASH cirrhotic liver tissue from Tzouanas et al (2026)^6^. The dataset comprised 9 control, 4 high-BMI MASH, and 5 low-BMI MASH tissue microarray generated using an in situ spatial transcriptomics platform. Cell-type annotations were derived from the original study.

### Statistical analysis

Differences in cluster prevalence and spatial organization across conditions were assessed using one-way ANOVA followed by Tukey’s post hoc test. Significance thresholds were defined as P < 0.05 unless otherwise specified.

### Gene enrichment and spatial differential analysis

ClusToRa includes an integrated gene enrichment analysis module for spatial differential analysis when molecular profiles are available. It first identifies statistically significant spatial interactions within each sample. Interactions were retained if they exceeded a user-defined enrichment threshold (default *Z* > 1.96), ensuring downstream analysis was restricted to robust colocalization events.

For each retained interaction, gene-level differential expression between colocalizing and non-colocalizing cells was assessed using a two-sided Wilcoxon rank-sum test. Effect sizes were computed as *log*_2_-transformed fold-changes with pseudocount stabilization:

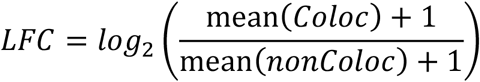

where *Coloc* denotes cells colocalizing with a given cluster and *nonColoc* denotes non-colocalizing cells. P-values were corrected for multiple hypothesis testing using false discovery rate (FDR) estimation (Storey procedure).

To identify conserved spatial transcriptional programs across samples of same group, interaction-level statistics were aggregated using a meta-analytic framework. For each gene, p-values from individual interactions were combined using Fisher’s method:

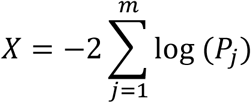

where *m* is the number of interactions and *P*_*j*_ denotes the interaction-specific p-value. The combined statistic follows a chi-square distribution with 2*m* degrees of freedom, and combined p-values were computed accordingly. To ensure robustness of aggregated signals, ClusToRa applies a directional consistency filter across interactions. For each gene, the proportion of interactions exhibiting concordant directionality (positive or negative *log*_2_ fold-changes) was computed:

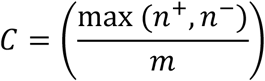

where *n*^+^and *n*^−^represent the number of positive and negative effects, respectively. Genes were retained for downstream analysis if *C* ≥0.75 (user adjustable), ensuring that aggregated signals reflect consistent spatial trends rather than mixed-direction effects. Group-level effect sizes were computed as the arithmetic mean of *log*_2_ fold-changes across interactions:

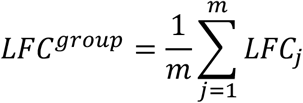

Finally, ClusToRa supports downstream functional analysis through gene enrichment using libraries such as Enrichr, enabling pathway-level interpretation of cluster-resident and colocalizing populations.

## Code and data availability

ClusToRa is available at: https://github.com/maringa780/ClusToRa. It can be executed as a standalone application for Windows and macOS, launched via MATLAB Online, or cloned for local use within MATLAB. The repository also includes an example workflow based on the Tzouanas et al. dataset, along with detailed documentation and illustrative videos demonstrating how to reproduce the analyses presented in this study.

## References

1. Longo, S. K., Guo, M. G., Ji, A. L. & Khavari, P. A. Integrating single-cell and spatial transcriptomics to elucidate intercellular tissue dynamics. Nat. Rev. Genet. 22, 627–644 (2021).

2. Dries, R. et al. Giotto: a toolbox for integrative analysis and visualization of spatial expression data. Genome Biol. 22, (2021).

3. Bull, J. A., Mulholland, E. J., Leedham, S. J. & Byrne, H. M. Extended correlation functions for spatial analysis of multiplex imaging data. Biological imaging 4, (2024).

4. Summers, H. D., Wills, J. W. & Rees, P. Spatial statistics is a comprehensive tool for quantifying cell neighbor relationships and biological processes via tissue image analysis. Cell Reports Methods 2, (2022).

5. Palla, G. et al. Squidpy: a scalable framework for spatial omics analysis. Nat. Methods 19, 171 (2022).

6. Tzouanas, C. N. et al. Hepatic adaptation to chronic metabolic stress primes tumorigenesis. Cell 189, 435–460.e28 (2026).

7. Devasia, A. G., Ramasamy, A. & Leo, C. H. Current Therapeutic Landscape for Metabolic Dysfunction-Associated Steatohepatitis. Int. J. Mol. Sci. 26, (2025).

8. Seubnooch, P., Montani, M., Dufour, J. F. & Masoodi, M. Spatial lipidomics reveals zone-specific hepatic lipid alteration and remodeling in metabolic dysfunction-associated steatohepatitis. J. Lipid Res. 65, (2024).

9. Li, Z. et al. Spatially resolved multi-omics of human metabolic dysfunction-associated steatotic liver disease. Nature Genetics 2025 57:12 57, 3112–3125 (2025).

10. Sasagawa, S. et al. Spatial profiling of chronic liver disease: a pilot spatial case series. Scientific Reports 2026 https://doi.org/10.1038/S41598-026-49400-7 (2026) doi:10.1038/S41598-026-49400-7.

11. Wen, W. et al. Integrating multi-omics and machine learning systematically deciphers cellular heterogeneity and fibrotic regulatory networks in the progression from MASLD to MASH. npj Digital Medicine 2026 9:1 9, 167– (2026).

12. Carter, J. K. & Friedman, S. L. Hepatic Stellate Cell-Immune Interactions in NASH. Front. Endocrinol. (Lausanne). 13, (2022).

13. Xie, D. et al. Interactions between hepatic stellate cells and immune cells: Implications for liver fibrosis. Biochim. Biophys. Acta Mol. Basis Dis. 1872, (2026).

14. Breuer, D. A. et al. CD8+ T cells regulate liver injury in obesity-related nonalcoholic fatty liver disease. 10.1152/ajpgi.00040.2019 318, G211–G224 (2020).

15. Kuleshov, M. V. et al. Enrichr: a comprehensive gene set enrichment analysis web server 2016 update. Nucleic Acids Res. 44, W90–W97 (2016).

